# Predicting the Conformational Variability of Oncogenic GTP-bound G12D Mutated KRas-4B Proteins at Cell Membranes

**DOI:** 10.1101/2021.07.19.452936

**Authors:** Huixia Lu, Jordi Martí

## Abstract

KRas proteins are the largest family of mutated Ras isoforms, participating in a wide variety of cancers. Due to their importance, large effort is being carried out on drug development by small-molecule inhibitors. However, understanding protein conformational variability remains a challenge in drug discovery. In the case of the Ras family, their multiple conformational states can affect the binding of potential drug inhibitors. To overcome this challenge, we propose a computational framework based on combined all-atom Molecular Dynamics and Metadynamics simulations able to accurately access conformational variants of the target protein. We tested the methodology using a G12D mutated GTP bound oncogenic KRas-4B protein located at the interface of a DOPC/DOPS/cholesterol model anionic cell membrane. Two main orientations of KRas-4B at the anionic membrane have been obtained and explored. The corresponding angles have been taken as reliable reaction coordinates so that free-energy landscapes have been obtained by well-tempered metadynamics simulations, revealing the local and global minima of KRas-4B binding to the cell membrane, unvealing reactive paths of the system between the two preferential orientations and highlighting opportunities for targeting the unique metastable states through the identification of druggable pockets.

## 1 Introduction

Cell plasma membranes (PM) are active agents in the regulation of protein structure and function, including mutant species involved in cancer diseases such as the proteins belonging to the Ras family ^1^. In particular, specific mutations in the family of Ras proteins are known to be responsible of their conversion to oncogenic species ^2,3^. Ras proteins are normally anchored at the cytoplasm and can be activated by guanosine-diphosphate (GDP)-guanosine-triphosphate (GTP) switching ^4,5^ following an incoming signal from their upstream regulators. The regulation between GDP and GTP is usually performed through the catalysis by GAP (GTPase Activating Proteins, like neurofibromin) or by activation of GEF (Guanine Exchange Factors, like the human regulator of chromosome condensation, RCC1). Since oncogenic Ras is defective in GAP-mediated GTP hydrolysis it results in an accumulation of constitutive GTP-bound RAS in cells. It has been observed that among the three mutated Ras isoforms in human cancers, KRas is the predominant one, found in 85% of the cases ^6^. In particular, it was found that KRas G12D mutation (instead of NRas G12D mutation) promoted colon cancer development in Apc-deficient mice, supporting the ability of KRas but not NRas to initiate the formation of such cancers ^7^, so that this especific mutation has been chosen to constitute the target oncogenic protein studied in the present work. Further, the variant KRas-4B was found at high concentrations in colorectal, pancreatic and lung cancer cells ^6,8,9^. Interestingly, it has been observed that Ras proteins are able to form nanoclusters of 6-7 units (also named proteo-lipid signaling complexes) with a radius size of about 9nm and able to survive in the time scale of 1 s ^10–12^. Such clusters can play a relevant role in the signaling of the protein. Recently, researchers have provided evidence that Ras dimerization may also be an important requirement for proper Ras activation of signaling pathways ^13–15^. Hence, controlling the orientations of the protein can influence the formation of Ras dimers, in such a way that signaling of Ras should be strongly affected.

KRas-4B can be divided into two distinguishable parts ^16^: a catalytic domain (CD) and a recently resolved ^17^ hypervariable region (HVR), formed by residues 167-185 of the protein, ending in a farnesylated (FAR) Cys-185 terminal. The CD is composed of residues 1-166, in a sequence which contains a catalytic lobe (the effector-binding domain, residues 1-86) and an allosteric lobe (residues 87-166). HVR preferentially binds the membrane in the liquid crystal phase, by inserting its farnesyl moiety into the phospholipid bilayer ^18,19^. Ras proteins are distinguished by their unique HVR post-translational modifications (PTM). The most important ones are acetylation, carboxymethylation, phosphorylation, prenylation and ubiquitination ^20,21^. For KRas-4B, the steps are given in a sequence (see Fig. 1): firstly the prenylation reaction, catalyzed by farnesyltrasferase (FTase) or geranylgeranyltransferase (GGTase), works by the addition of an isoprenyl group to the Cys-185 side chain. Then farnesylated KRas-4B is hydrolyzed and catalyzed by the endopeptidase enzyme called Ras-converting enzyme 1 (RCE1). In this step the VIM motif (HVR tail composed of three amino acids: valine-isoleucine-methionine) of the C-terminal Cys-185 is removed. Later on, KRas-4B is carboxymethylated at the carboxyl terminus of Cys-185 catalyzed by isoprenylcysteine carboxyl methyltransferase, forming a reversible ester bond ^22^. As shown in Fig. 1, this reversible reaction can modulate the equilibrium of methylated KRas-4B (KRas-4B-FMe) and demethylated KRas-4B (Kras-4B-Far) population in tumors with potential huge impact on downstream signaling, protein-protein interactions and protein-lipid interactions ^23^.

**Fig. 1.**
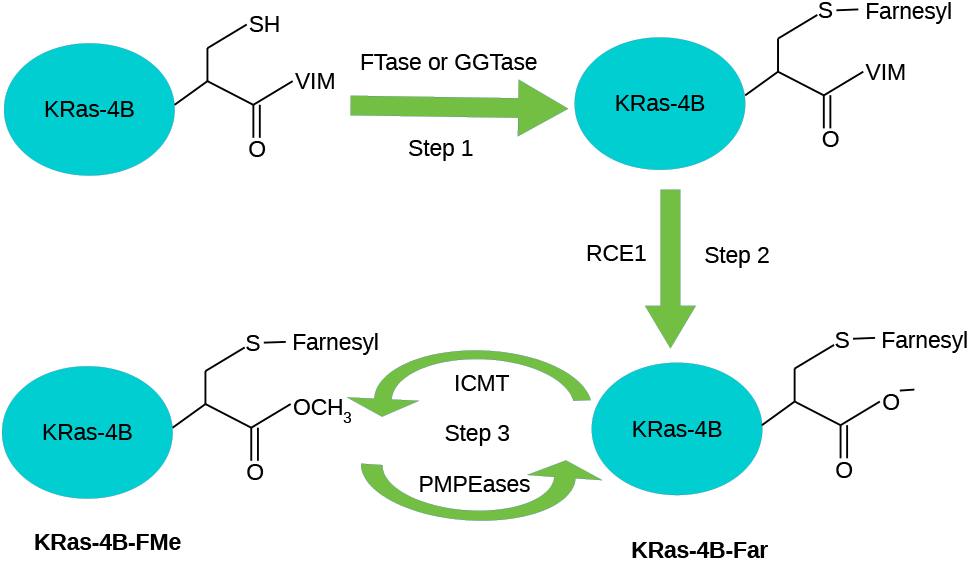
PTMs steps of KRas-4B in cells: prenylation, hydrolysis, carboxymethylation and decarboxymethylation.

It has been recently found that the two major pathways in oncogenic Ras-driven proliferation can be promoted in the case that KRas is membrane-anchored ^24^. The knowledge of structural details on GTP-bound KRas-4B will help to design oncogenic KRas-4B inhibitors ^25^. In this way, methods of disabing interactions between KRas-4B and the cell membrane can be devised and become a crucial help to target anticancer drug discovery ^20^. The translocation pathways of Ras proteins vary for different Ras isoforms, and signaling activity of KRas-4B is dependent on its enrichment level in the PM. Phosphodiesterase*δ* (PDE*δ*) has been revealed to promote effective KRas-4B signaling by sequestering KRas-4B from the cytosol by binding the prenylated HVR and help to enhance its diffusion to the PM throughout the cell, where it is released to activate various signaling pathways required for the initiation and maintenance of cancer ^17,26^. This is one of the common targets considered for oncological drug development. Research of the mechanistic inhibitory treatment of PDE*δ* has been carried out ^26,28^ with the aim to identify a panel of PDE*δ* inhibitors. It has been reported ^17^ that the affinity between KRas-4B-FMe and PDE*δ* is 78-fold compared with its fanesylated demethylated variant by using isothermal titration calorimetry measurements. Further, a 5-amino-acid-long sequence motif in its HVR (K-S-K-T-K), which is shared by both methylated and demethylated KRas-4B, may enable PDE*δ* to bind KRas-4B-FMe. It has also been observed that there is a clear difference in the level of the reversible carboxymethylation of Cys-185 according to different kinds of tumor types related to KRas-4B ^27,29^. As it was described in a very recent review ^30^, the use of computational modeling and simulation to achieve a deeper understanding of Ras signaling is expected to allow us the design of new mechanism-based therapies for Ras cancers.

In a recent work a detailed study of the demethylated KRas-4B-Far was reported ^19^. It was observed that strong, long-lasting salt-bridges between the FAR and HVR moieities are able to stabilize the binding of the oncogenic KRas-4B-Far to the PM for long periods of time, of the order of microseconds. Further, free-energy barriers along with the corresponding free-energy landscape (FEL) for KRas-4B-Far binding with the anionic model membrane were described. The main orientational angles of KRas-4B-Far when bound to the cell membrane in a fully quantative fashion ^31^ have been recently reported. In the present work, our aim has been to investigate the alternative farnesylated and methylated KRas-4B species (KRas-4B-FMe) and its structural, orientational and energetic characteristics. We have employed all-atom molecular dynamics (MD) and the FEL have been obtained by and well-tempered metadynamics (WTM) simulations ^19^, in order to acquire precise information on the orientation of the full-length KRas-4B and its preferential conformations when binding with the anionic bilayers. The proposed angular variables will be employed as collective variables (i.e. reaction coordinates) in the WTM simulations, so that the corresponding free-energy landscape will be able to reveal new insights into the underlying biological mechanisms, help to identify druggable pockets and foster molecular targeting ^32^, offering new opportunities in the design and discovery of new drugs able to deactivate the oncogenic species.

## 2 Methods

### 2.1 System Preparation and MD Simulations

We conducted 1 *μ*s long MD simulations of a single KRas-4B inside model anionic cell membranes constituted by DOPC (56%), DOPS (14%) and cholesterol (30%). MD is a poweful tool based in the generation of Newtonian trajectories for each atom in the system and able to handle from simple biological solvents (water bulk in a wide variety of thermodynamic states ^33–36^ or in solution ^35,37^) to complex ones, such as proteins ^44–47^ or membranes ^48–51^) and also able to render microscopic properties connected directly to experiments, such as all sorts of neutron diffraction and spectroscopic data (X-ray, infrared and Raman spectroscopy, etc) ^52,53^ to mention just a few examples. The system contains the KRas protein and a total of 304 lipid molecules fully solvated by 60,000 TIP3P water molecules ^54^ in potassium chloride solution at the human body concentration (0.15 M), yielding a system size of 222,000 atoms. The temperature was fixed at 310.15 K and the pressure was set ot 1 atm, well above the corresponding transition temperature for DOPC and DOPS, i.e. at the liquid crystal state.

All MD inputs were generated using CHARMM-GUI membrane builder ^55,56^ and the CHARMM36m force field ^57^ was adopted for lipid-lipid and lipid-protein interactions. Crystal structure of KRas-4B with partially disordered hypervariable region (pdb 5TB5) and GTP (pdb 5VQ2) were used to generate full length GTP-bound methylated KRas-4B proteins. Both pdb files were downloaded from RCSB PDB Protein Data Bank ^58^. The full length KRas-4B-FMe and GTP were solvated in a water box and equilibrated for 20 ns before generating the full setups. KRas-4B-FMe was initially set anchoring to membrane at the beginning of the simulations. Additional techincal details of the MD simulations are reported in “Supporting Information” (SI).

### 2.2 Well-tempered Metadynamics

After equilibrating properly the system composed by oncogenic KRas-4B-FMe and the model cell membrane from the previous MD 1 *μ*s run, we switched to run another 1 *μ*s of WTM simulations to perform Gibbs free-energy calculations of KRas-4B-FMe binding at anionic phospholipid membrane bilayers, starting from the last configuration of MD simulations. The problem of computing FEL in multidimensional systems has been extensively discussed for decades. Among the variety of methods proposed, we can highlight transition path sampling ^59,60^, which does not require the knowledge of reaction coordinates and it has successfuly been applied to simplified lipid bilayers ^61–64^ or methods where the relevant coordinates must be previously defined, such as adaptive biasing force ^65^, multistate empirical valence bonds ^5,66,67^ and umbrella sampling ^68,69^, among the most popular. In the present work we have employed WTM because of its suitability for a wide variety of systems, including model cell membrane systems with attached small-molecules and proteins ^19,70–76^. Some especific details of the method will be given below and the main parameters of WTM simulations are listed in SI.

1 *μ*s long well-tempered metadynamics simulations were performed using the joint GROMACS/2018.3-plumed tool ^77^. The isothermal-isobaric NPT ensemble at temperature 310.15 K and pressure 1 atm was adopted in all cases. Periodic boundary conditions in the three directions of space were also considered. The two collective variables (CV), which were adopted in the WTM simulations are shown in Fig. 2. They were considered to describe the conformational changes along free-energy paths, as recently described in a study devoted to KRas-4B-Far proteins anchoring to the anionic model membranes with different cholesterol concentrations ^31^.

**Fig. 2.**
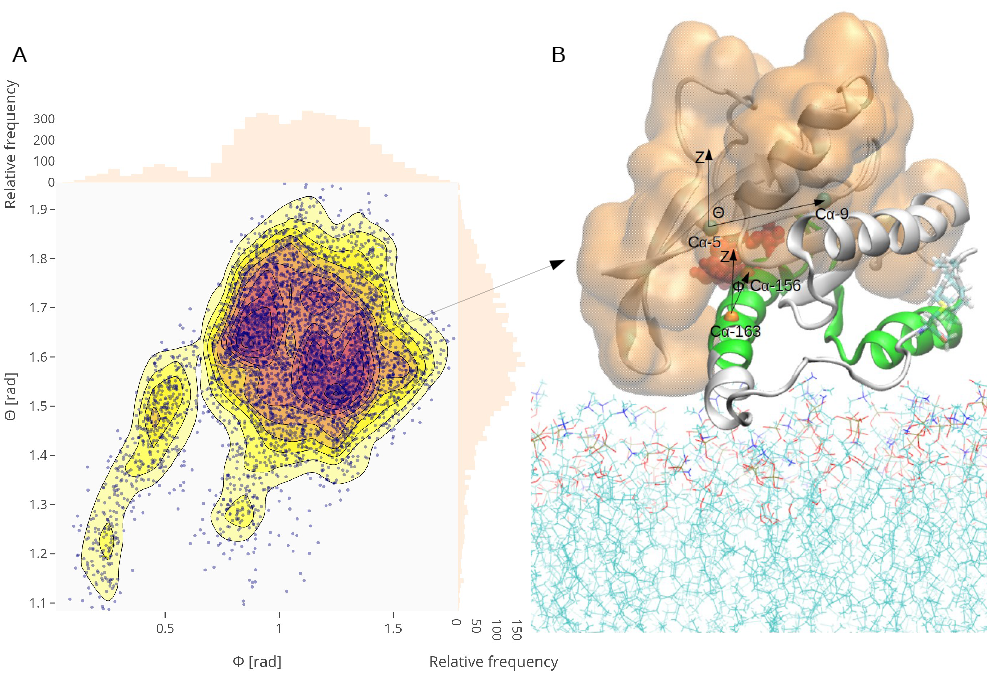
Orientation of the mutant KRas-4B-FMe on the anionic membrane. Panel A shows the density distribution of conformations projected onto a plane defined by the reaction coordinates Φ and Θ (rad). The relative frequency of each coordinate is shown on the right and upsides. Panel B displays a typical conformation of the G12D mutated and Ser-181 phosphorylated KRas-4B-FMe binding with the anionic membrane. CV that will be adopted in the WTM simulations are shown here. The putative dimerization interface (*α*4-*β* 6-*α*5 region) of KRas-4B-FMe is shown in green. Lipids in the membrane are shown as thin lines. The effector-binding domain is highlighted in orange. For the sake of clarity water and ions are not shown.

## 3 Results

As indicated above, in the present work we have considered the KRas-4B-FMe protein, resulting of the mutation of the wild-type KRas-4B at Gly-12 aminoacid (G12D mutation), phosphorylated at the Ser-181 site (PHOS site), and further methylated at the FAR site. Sketches of parts of the full system are described in Fig. 1 of SI.

In order to explore structural characteristics of the anchoring of KRas-4B-FMe proteins at anionic membranes, several physical properties of the membrane have been calculated. The area per lipid, thickness of the bilayer, and the deuterium order parameter *S_CD_* of this system have been reported in a previous work ^19^, confirming a good test for such validation of MD simulations of cell membranes. In the present study, we obtained an averaged area per lipid of 0.524 ± 0.006 nm^2^, very close to the 0.544 nm^2^ obtained by Nagle et al. ^78^ for a DOPC:cholesterol (30%) bilayer by means of X-ray scattering in the low and wide angle regions at 303 K. The thickness (Δ*z*) of the membrane has also been computed as the mean distance between phosphorus atoms of the DOPC head groups from both leaflets. We have obtained a value of 4.34 ± 0.05 nm, in overall good agreement with the experimental value of Δ*z* = 3.94 nm for a pure DOPC/DOPS (4:1) bilayer at 303 K, as reported by Novakova et al. ^79^. We should point out that the presence of cholesterol produces a remarkable reduction of the area per lipid and a clear increase of the thickness of the membrane because of the more significant packing of the surfactant molecules ^80,81^.

In the case of demethylated KRas-4B-Far species, we previously reported that active hydrogen (−NH_3_) and oxygen atoms from the side chains of HVR and CD can establish stable hydrogen bonds (HB) called salt-bridges with the interface of the membrane (see Fig. S7 and S8 of Ref. ^19^) which help to stabilize KRas-4B-Far binding with the interface of the membrane, regardless of its farnesyl group anchoring into the membrane or not. In the present case, we have observed that under the circumstances of phosphorylation at Ser-181, the O _*f me*_ of farnesyl group forms typical HB with the phenoxyl hydrogen atom belonging to Tyr137 of CD (see Fig. 2 of SI) while salt-bridges exist between the hydrogen atoms of the cationic ammonium 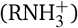 from the lysine of HVR of KRas-4B-FMe and the anionic oxygen atoms from phosphorylated Ser181 (see Fig. 2 of SI), resulting in the departure of the farnesyl group of KRas-4B-FMe.

### 3.1 Preferential conformations of G12D mutated KRas-4B-FMe on the model membrane

Given the central importance of the proper orientations of KRas proteins on cell membranes to the signal transductions in or between cells, orientations of the CD of KRas-4B-FMe are displayed considering the last 975 ns of the MD trajectory. We have calculated the density distribution of conformations of KRas-4B-FMe projected onto a plane defined by two specific angular, torsional variables Φ and Θ during the last 500 ns simulation time and the result is presented in Fig. 2. The torsion angle Φ is defined by the membrane normal and a vector running the C_*α*_ atoms of residues 163 and 156 on the last helix *α*5 of lobe 2, whereas the torsion angle Θ is defined between the membrane normal direction and a vector running the C_*α*_ atoms of the residue 5 and the residue 9 which belong to the first strand *β* 1 in the structure of KRas-4B ^82^.

From Fig. 2, we can observe that the mutant KRas-4B-FMe spends most of the time in the active state centered around (1.0 rad, 1.6 rad), exposing the effector-binding domain for further signal transduction. Obviously, in the last 500 ns trajectories of MD simulations, the mutated KRas-4B-FMe is trapped in the active state because of the strong interactions between HVR and the interface of the bilayer, ^19^ and seldom does KRas-4B-FMe explore other conformations blocked by high energy barriers. We continued to explore the corresponding free-energy landscape by conducting WTM simulations. By doing so, energy barriers on the potential energy surface might be overcome, allowing for the exploration of new conformational space.

### 3.2 Well-tempered metadynamics simulations and conformational transitions of KRas-4B: Two-dimensional free-energy landscapes

The enhanced sampling method WTM was used to analyze the conformational transitions of KRas-4B-FMe. Metadynamics applies a time-dependent biasing potential along a set of CV by adding Gaussians to the total potential in order to overcome barriers larger than *k_B_T*, where *k_B_* is Boltzmann’s constant and *T* is the temperature. In the method, sampling is facilitated with an additional bias potential which acts on a selected number of degrees of freedom (*s_i_*), the so-called CV along the trajectory. For each degree of freedom the biased potential is constructed as the sum of Gaussian functions ^83–85^:

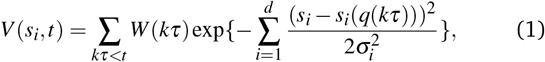

where *k* is an integer, *τ* is the Gaussian deposition stride, *W* (*kτ*) is the height of the Gaussian and σ_*i*_ is the width of the Gaussian for the i-th CV. The choice of CV is extremely important. A good CV must be able to distinguish between the initial and final states while properly describing all relevant intermediate states. The CV usually considered are relative distances between atomic sites of the protein (and eventually GTP) and the center of the membrane (see for instance ^19^) able to indicate the main positions of the target molecule at the most relevant stable states. In the present work, given the shorter durability and strenght of the interactions of FAR and the cell membrane, we have taken a different approach to analyze the free-energy landscape of KRas-4B-FMe. Following studies showing the significant influence of orientations of Ras proteins on membranes related to their function in cell and considering that cell membranes are platforms for cellular signal transduction ^82,86^ we reported in a recent study ^31^ two preferential orientations of oncogenic and wild-type KRas-4B-Far proteins able to describe the reorientations of the protein while anchored at the membrane. Henceforth, we considered the two particular orientations, Θ and Φ defined above as the CV of the WTM method, we have run 975 ns WTM simulations and obtained the corresponding free-energy hypersurface.

Technical details and a full study of the convergence of the simulations are reported in SI. The two dimensional free-energy landscape is represented in Fig. 3 for the two selected CV. We can easily observe that two main basins (A, B) are well defined so that a main transition state (TS) along the MFEP arises around coordinates (1.51,1.51), as indicated in Table 3 of SI, although other secondary basins (C, D, E) and lower TS can be also observed. The energy barrier of such TS between A and B will indicate the amount of free-energy required for KRas-4B-FMe to exchange its conformation between the two principal orientations, as it was described in Ref. ^31^. The two stable states A and B are represented in Fig.4: there we can observe how the main structure of the CD at state A is somehow farther away from the interface and the center of the membrane, with the HVR in close contact with the interface, whereas in state B, the protein has suffererd a rotation putting the CD deeper at the interface but hidding HVR and FAR group inside the main body of the protein.

**Fig. 3.**
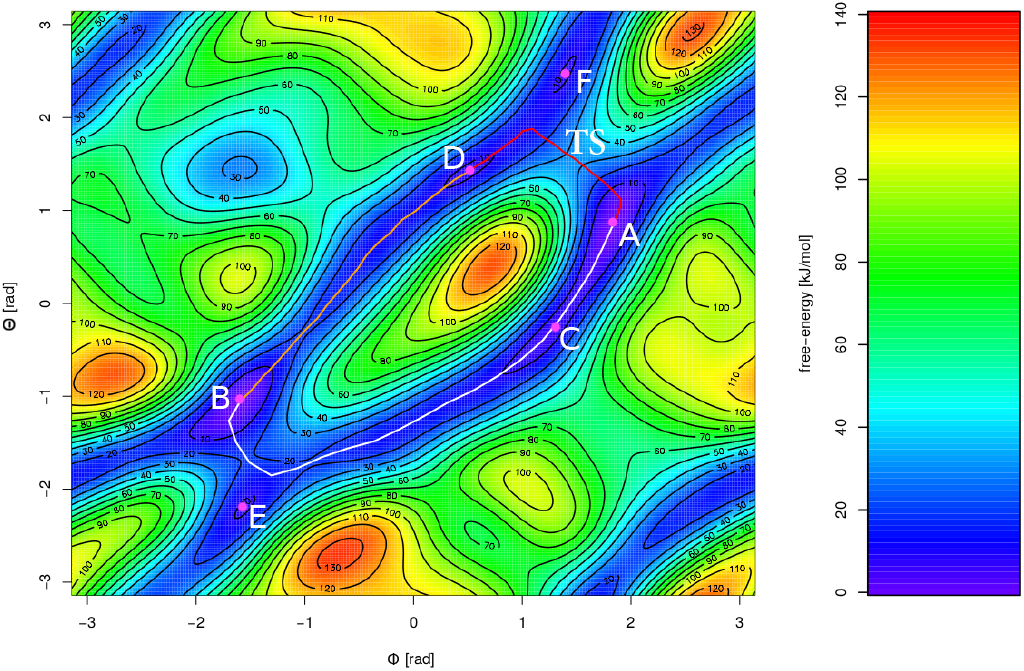
2D free-energy landscape F(Φ, Θ) (kJ/mol) for the oncogenic KRas-4B-FMe. Basins ‘A’, ‘B’, ‘C’, ‘D’, ‘E’, and ‘F’ are the (meta-)stable states and are depicted in pink color. A single MFEP is depicted in different colors between different basins. ‘TS’ indicates the transition state between ‘A’ and ‘D’.

**Fig. 4.**
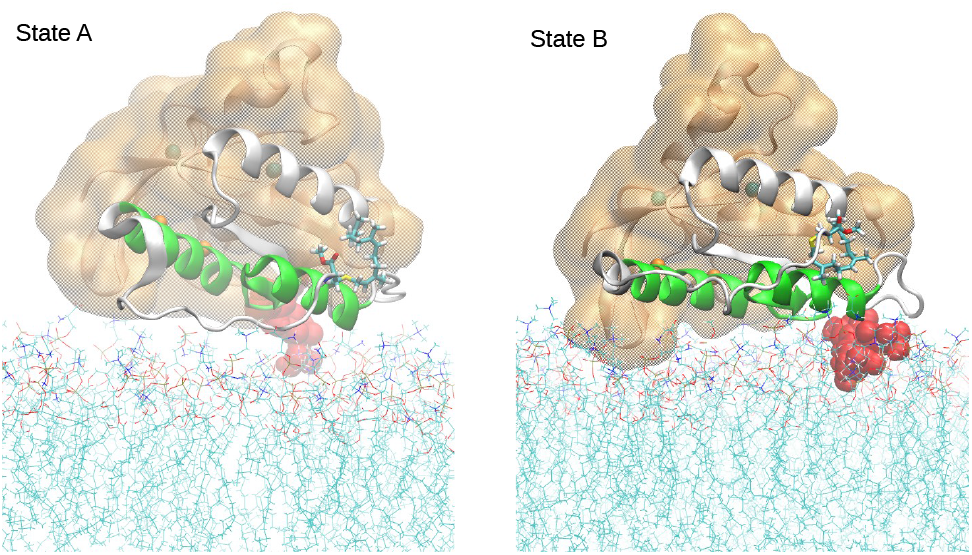
Configurations of KRas-4B-FMe in Basins ‘A’ and ‘B’. Colors defined as in Fig. 2. GTP is represented in red.

Using standard procedures, we have been able to trace the minimum free-energy path (MFEP) between the two stable states with high accuracy. MFEP can be traced by iteratively refining a pathway connecting stable states that converges to the minimum free-energy trajectories between them. MFEPs also allow us the determination of transition states. By means of the guidelines described in SI, the reconstruction of the obtained 2D free-energy surface was performed with R plugin metadynminer ^90^. Metadynminer was also used to identify the MFEP along the free-energy landscape and eventually locate TS. The coordinates of the minima of the FEL are reported in Table 1 and the corresponding main TS between the two stable states is indicated in Table 3 of SI.

**Table 1.** Minima on the FEL of the mutated KRas-4B-FMe binding with the anionic membrane. Here each letter refers to one (meta-)stable state, among which A stands for the global state which is set to zero. Locations of minima of FEL of Fig. 3 were obtained by using metadynminer package.

| States | $\Phi$ (rad) | $\Theta$ (rad) | Free-energy (kJ/mol) |
| --- | --- | --- | --- |
| A | 1.78 | 0.77 | 0.00 |
| B | -1.58 | -1.04 | 0.84 |
| C | 1.19 | -0.38 | 5.14 |
| D | 0.48 | 1.39 | 6.34 |
| E | -1.59 | -2.25 | 7.59 |
| F | 1.49 | 2.67 | 8.18 |

The free-energy barrier from basin B to basin A has been estimated to be of 18.39 kJ/mol along the MFEP depicted in Fig. 3, whereas ΔF from ‘A’ to ‘D’ = 30.98 kJ/mol and ΔF from ‘D’ to ‘B’ = 11.07 kJ/mol, all along the MFEP.

In order to establish a connection between the two preferential orientations of the methylated KRas-4B and the exposing docking pockets to be targeted by anti-cancer drugs, we have considered the C*α* root-mean-square difference (RMSD) of the CD with respect to the crystal structure of KRas4B (see Ref. ^19^) from the standard MD simulations. The results are shown in the upper panel of Fig. 5. We show how does the structure change along with the simulation. Values of RMSD around 3-5 Å or less normally indicate a well equilibrated MD simulation, as is the case in the present work. Further, we can consider the so-called root-mean-square fluctuation profile (RMSF), shown in the bottom panel of Fig. 5. We only considered the C*α* of every residue to generate the RMSF. Usually, regions of KRas-4B that can be targeted by therapeutic drugs lie in Switch-I or Switch-II pockets (see Ref. ^91^). The location of the Switch-I/II pockets is indicated in Fig. 1 of Ref. ^24^. (Switch I: residues 30-38 and Switch II: residues 60-76). In Fig. 5, we can see that these two drugging pockets that locate in the effector-binding domain (residues 1-86), as well as the HVR tail, have higher structural flexibility. Apparently, the motion of the allosteric lobe (residues 87-166) of KRas-4B-FMe is limited by its rigid structure and contributes much less to the RMSD profile than the effector-binding domain of KRas-4B-FMe. In our MD study, we observe that Switch I and the HVR region are the areas suffering larger vibrational ranges, with amplitude of 1-2 Å. In the subsequent WTM work, we have designed the two CV to focus on how do structural changes influence the free-energy profile of KRas-4B-FMe along with the corresponding CV space (see Fig. 3). In our characterization, the transition between basins ‘A’ and ‘B’ mainly corresponds to the vibrations of the proper druggable pockets (Switch I/II) pockets observed in Fig. 5.

**Fig. 5.**
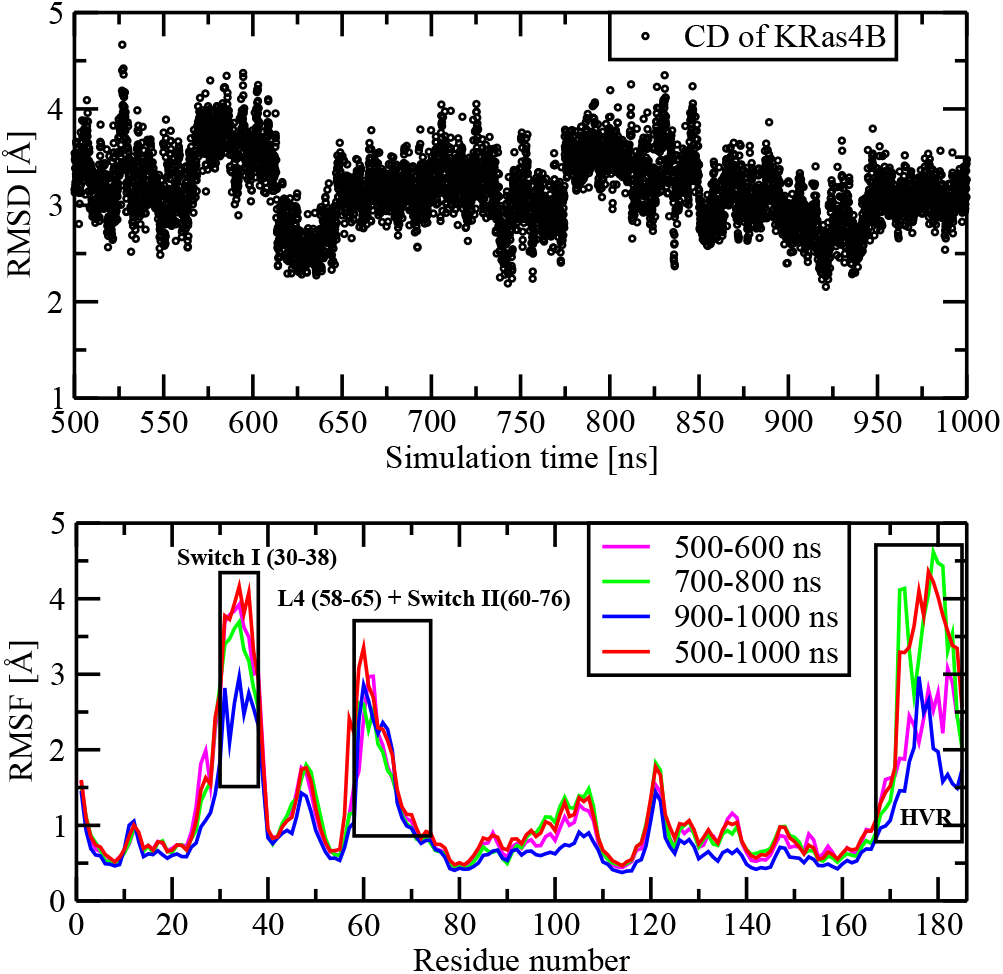
Root mean-square difference of the catalytic domain of KRas-4B and root mean-square fluctuation profiles. The latter indicates the time evolution of highlighted residues along the second 500 ns period of the MD simulation.

### 3.3 One-dimensional free-energy profiles

From the two-dimensional free-energy landscape obtained from WTM simulation (Fig. 6) is also possible to obtain a one-dimensional (1D) free-energy profile where one single CV is considered and the second one has been integrated out. This calculation along a single CV (*s_i_, i* = 1, 2), allows us to directly compare free-energy barriers to experimental findings ^92^. 1D free-energy profiles F(*s*_1_) can be obtained as follows ^92,93^:

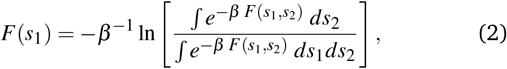

where *s*_1_ and *s*_2_ are the CV, *β* = 1*/*(*k*_B_*T*), *k*_B_ is the Boltzmann constant and T is the absolute temperature. This means that all possible paths for the CV labelled as *s*_2_ have been integrated out and averaged. *F*(*s*_1_) reveals additional information about stable states and free-energy barriers only as a function of one single CV. In the present case the results reveal quasi-symmetrical profiles for Θ and some asymmetry for Φ, indicating the existence of two equivalent orientations for each coordinate. However, the barriers related to the change from one orientation to the second are different in size, of about 15 kJ/mol for Φ and of 8 kJ/mol for Θ. These values are much lower than the global barrier reported above which corresponds to the joint reorientation of the two torsion angles. Interestingly, Zhou et al. ^4^ reported results of metadynamics simulations of K-Ras in plasma membranes on the conformational changes also using angular collective variables: CV1 was the C*α*-atom RMS of residue 177-182 from a helical structure and CV2 the one end-to-end distance involving the C*α*-atom of residue 176 and 184, at the 1 *μ*s timescale. These authors found barriers of the order of 12 kJ/mol, quite close to the ones reported in the present work.

**Fig. 6.**
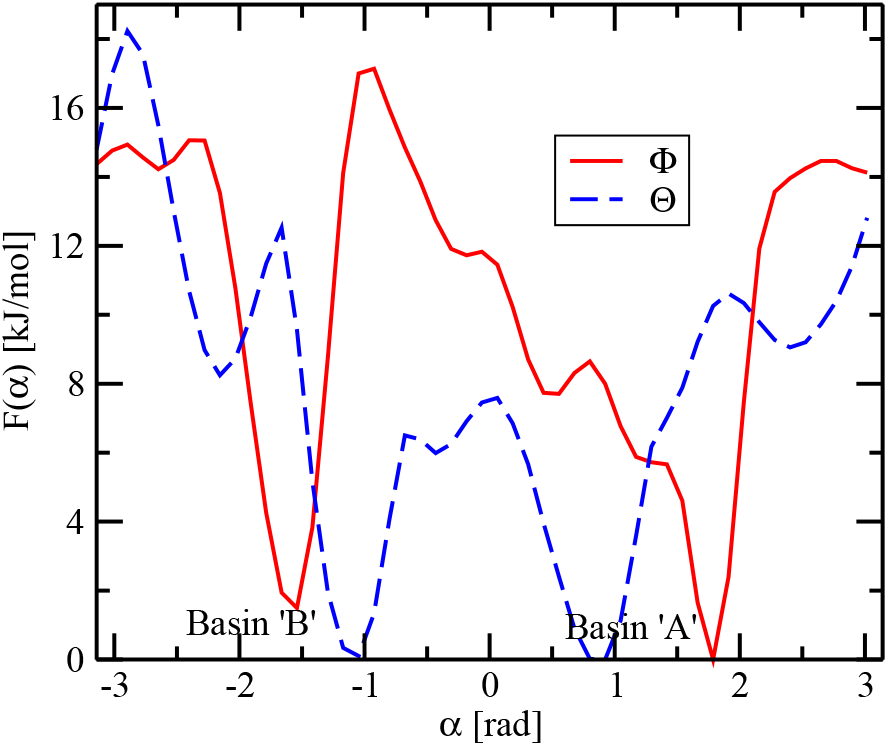
1D integrated free-energy profiles for the two CV (*α* = Θ, Φ.) Basins marked with same labels as in Fig. 3. Minima are set equal to zero. Free-energy profiles generated by the PLUMED2 tool.

## 4 Conclusions

KRas-4B are small GTPase with the ability to regulate cell growth, differentiation and survival and are related to cancers such as in lung, colon and pancreas. Even though great efforts have been focused on the methylated KRas-4B protein, fundamental aspects such as its main conformations, bonding sites to the cell membranes and its free-energy landscape remain very much unknown.

In this work, we have conducted MD and WTM simulations of a GTP-bound oncogenic KRas-4B-FMe protein bound to a model cell membrane composed by DOPC (56%), DOPS (14%) and cholesterol (30%) inside KCl solution at 310.15 K and at the fixed pressure of 1 atm. Our simulations reached the scale of 1 *μ*s. The oncogenic KRas-4B protein contains two mutations: G12D and phosphorylation at its Ser-181 site in addition to the methylation of its farnesyl terminal.

We have characterised the system by computing area per lipid and thickness of the membrane and corroborated their good overal agreement with experimental data. In a previous work ^19^, we observed the existence of long-lasting salt bridges between farnesyl and the phosphorilated site of the HVR part of the protein able to stabilize the anchoring of the oncogenic protein to the membrane. In the present case of the methylated protein, these bonds still exists but they are much weaker. According to our results, the anchoring of the KRas-4B species to the membrane in the present case is essentially due to hydrogen bonding between HVR and the phosphoserine group of the protein and also by oxygen sites of the farnesyl group and hydrogens pertaining to the tryosine sites of the CD (see Fig. 2 of SI). So, the information reported in this work could be of help for the development of farnesyltransferase inhibitors acting as anti-cancer agents ^94^.

The free-energy profiles of conformational orientations of the mutated and methylated protein computed by WTM have given us quantitative information on the energy barriers that must be surmounted by KRas-4B to change its orientation. We considered as CV of the method two angles previously defined ^31^, able to clearly distinguish variations of the orientation of the protein. These two angles (Φ, Θ) are able to describe two classes of angular reorientations: one related to changes of one single coordinate and a more complex one corresponding to joint changes of the two CVs. With the help of well-tempered metadynamics simulations we have calculated 2D FEL and the 1D free-energy profiles along one CV. We have two global stable states of KRas-4B linking to the membrane and a transition state between them. Our results indicate that for GTP-bound oncogenic KRas-4B-FMe there exist free-energy barriers of 20 or 30 kJ/mol that need to be surmounted by the protein in order to shift between one stable state to the other. This is a large barrier and it suggests that when KRas-4B-FMe is attached to the membrane in a particular orientation, i.e. in a specific pair of torsinal angles, it will require long periods of time and eventual increase of temperature or pressure to be able to access its second preferential stable orientation. From the calculation of 1D integrated free-energy profiles we have observed that reorientations along one single angular variable are much common, since the corresponding barriers are only of the order of 10 kJ/mol. The preliminary analysis of the two main drugging pockets of KRas-4B indicate that the preferential orientations reported in the present work should be associated with the Switch I/II pockets as reported in Refs. ^24,91^.

## 5 Tables

## Acknowledgements

We thank financial support provided by the Spanish Ministry of Science, Innovation and Universities (project number PGC2018-099277-B-C21, funds MCIU/AEI/FEDER, UE).

