## Supporting Information for "Predicting the Conformational Variability of Oncogenic GTP-bound G12D Mutated KRas-4B Proteins at Cell Membranes"

Journal Name

ARTICLE TYPE

Cite this: DOI: 00.0000/xxxxxxxxxx

Huixia Lu[a] and Jordi Martí[*b]

#### 1 Characteristics of the Molecular Dynamics simulations

The components of the system considered in this work are reported in Fig. 1. Species GTP, DOPC, DOPS and cholesterol are sketched. The sequence of aminoacids forming KRas-4B is indicated and the detailed structure of the farnesyl tail is also shown, including the methyl group introduced to KRas in the methylation process. The sites indicated in red correspond to relevant species that will be referred to in the text and the figures. The component aminoacids of KRas-4B are listed in Table 1.

The system was first energy minimized and then equilibrated in Molecular Dynamics (MD) runs of 10 ns. Production runs were performed within the NPT ensemble for 1 $\mu$s. The GROMACS/2018.3 package was employed[1] for the MD simulations. Time steps of 2 fs were used in all production simulations and the particle mesh Ewald method with Coulomb radius of 1.2 nm was employed to compute long ranged electrostatic interactions. The cutoff for Lennard-Jones interactions was set to 1.2 nm. Pressure was controlled by a Parrinello-Rahman piston with damping coefficient of 5 $ps^{-1}$ whereas temperature was controlled by a Nosé-Hoover thermostat with a damping coefficient of 1 $ps^{-1}$. Finally, periodic boundary conditions in three directions of space have been taken.

As a complementary information, we report here in Fig. 2 two selected radial distribution functions (RDF) for selected sites in the hypervariable chain (HVR) and the catalitic domain (CD) (see Fig. S5 of Ref.[2], where hydrogen of lysine group belongs to HVR and hydrogen of the tyrosine group belongs to CD) correlated with hydrogen sites in the farnesyl (see Fig. 1) and phosphoserine (see Fig. S5 of Ref.[2]) groups. In both cases, the existence of hydrogen bonds between hydrogen and oxygen species are revealed, with typical distances of 1.7 Å for HVR-phosphoserine and of 1.85 Å for farnesyl-CD.

[a‡]School of Pharmacy, Shanghai Jiaotong University, Shanghai,China.
[b‡]Department of Physics, Technical University of Catalonia-Barcelona Tech, B5-209 Northern Campus, Jordi Girona 1-3, 08034 Barcelona, Catalonia, Spain.
* Corresponding author
‡These authors contributed equally to this work

#### 2 Convergence of the well-tempered metadynamics simulations

The values for the parameters of the well-tempered metadynamics (WTM) simulations[3] are listed in Table 2. The convergence of WTM simulations needs to be assessed because it is important to ensure that the number of transition events between stable states is statistically meaningful and also that sampling of all phase space of the collective variables (CV) has been achieved[4]. In the case of protein-bilayer systems with hundreds of thousands of atoms, to cover the full range of the CV space is a challenge. In the present work, two angular CV ($\Phi$ and $\Theta$ as defined in Section 3 of the manuscript) were selected.

Usually the size of the hills of the Gaussian kernels deposited along the simulation is monitorized. As the simulation progresses and the added bias grows, the Gaussian height would be progressively reduced, eventually including low-height spikes. In the present case, the height of the biased potential decreased accordingly along the simulation run, as indicated in Fig. 3. A quasi-flat profile is already seen after 450 ns, although some spikes due to large fluctuations in the values of CV are observed.

Despite the fact that the Gaussian height is decreasing to zero, it has been extensively shown that other measures of convergence of a metadynamics simulation must be explored in order to make sure the system is fully converged. To do this, we will proceed as follows:

1. The time evolution of the two CV along the full time span of our simulations has been reported in Fig. 4, we can see that the system is diffusing efficiently in the collective variable space and the bias potential produces that both CV are able to diffuse in full phase space. We can only observe a few regions where the diffusion was less efficient, indicating angular configurations not likely to reach and corresponding to the regions of highest free-energy as shown in Fig.3 of the main text.

2. To assess the convergence of the WTM simulation we have primarily calculated the integrated free-energy profiles as a function of simulation time (see Fig. 5). As we can see that the estimated free-energies do not change significantly in the final part of our simulations (starting at 550 ns) which also

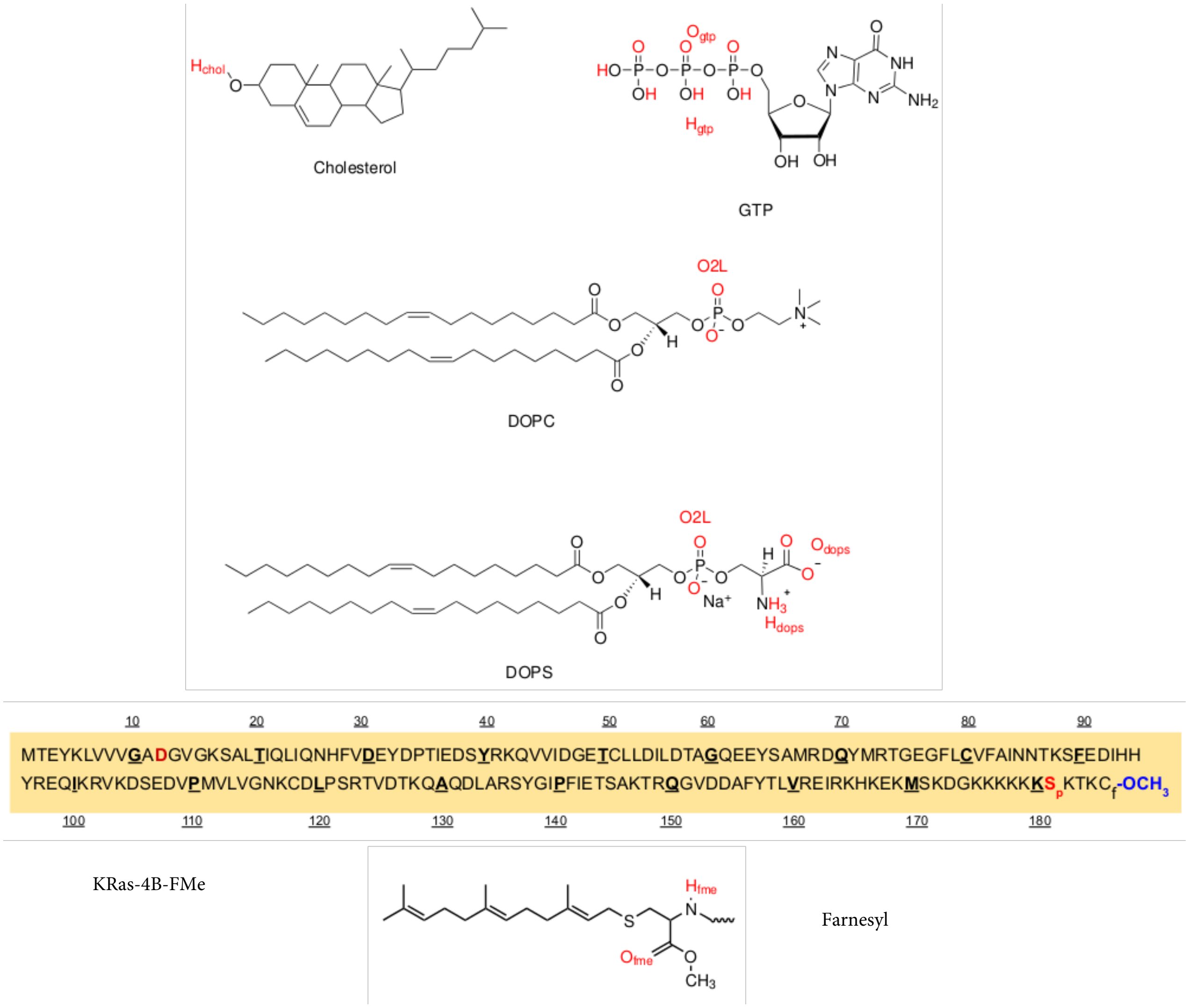


**Fig. 1** Sketches of GTP, DOPC, DOPS, cholesterol, KRas-4B-FMe, with details of the farnesyl (FAR) group, labeled as $C_f$.

ensures us the convergence of the simulations.

3. Convergence can be further assesed in different ways with more precision. We have considered two additional methods: (1) reporting the *block analysis* of the average error along free-energy profiles as a function of the block length and (2) performing the *committor analysis* of the CV (see PLUMED project[5,6]). In method (1) (see Fig.6) we report the size of the average error in the free-energy profiles calculated from two different sets of well-tempered metadynamics simulations as a function of the block size. The initial 25% of all metadynamics trajectories were discarded. As expected, the errors increase with the block length until they reach a plateau in all cases. The average errors tend to stabilise around 0.42-0.43 kJ/mol for the two considered CV. For this calculation, scripts provided by the PLUMED project[5,6] have been employed. Method (2) consists in throwing a large number of short MD simulations started at the local transition state (TS) between the two basins (A, B, as indicated in Figs. 3 and 4 of main text) and check whether the final values of those CV end up in one of the two stable basins of the free-energy hypersurface. After throwing around 500 MD trajectories started at the TS, we found (see Fig. 7) that 47.6 % of them ended in basin A and 52.4% ended in basin B. Given the similar probabilities we can conclude that the choice of CV was successful.

#### 3 Calculation of the minimum free energy paths

In order to obtain and evaluate the paths connecting the two stable states located on the 2D free energy landscape we have considered a process including three steps: (a) fixing the two local minima; (b) locating a coarse path and (c) refining the path. The coordinates of the path are numerically reported in Table 3. The first coordinate of each point corresponds to CV1 ($\Phi$) and the final coordinate to CV2 ($\Theta$). The R-package metadynminer[7] reads HILLS files from PLUMED, calculates free energy surface by fast Bias Sum algorithm, finds minima and analyses transition paths by Nudged Elastic Band method.

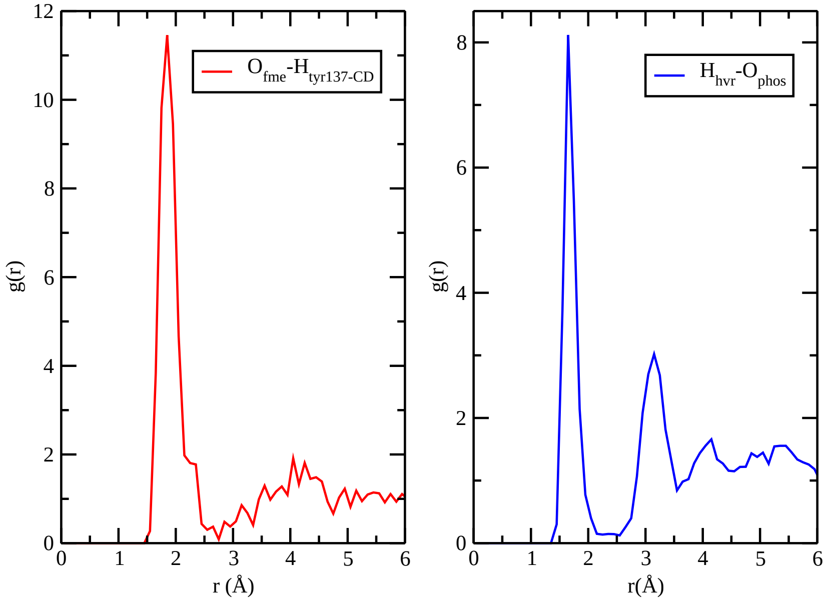


**Fig. 2** Selected RDF for active hydrogens of HVR (right) and CD (left) with selected sites of FAR ($O_{fme}$) and phosphoserine group ($O_{phos}$) of the KRas-FMe.

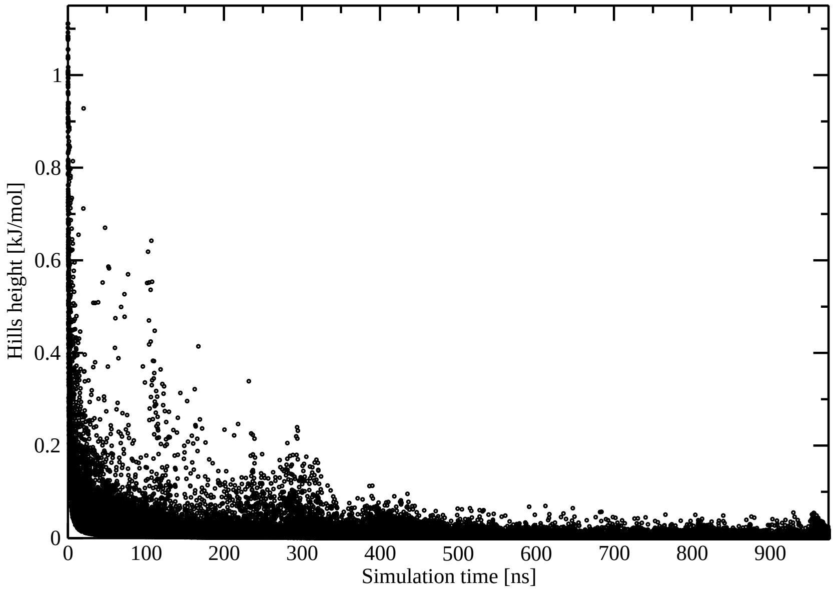


**Fig. 3** Well-tempered Metadynamics hills height as a function of simulation time.

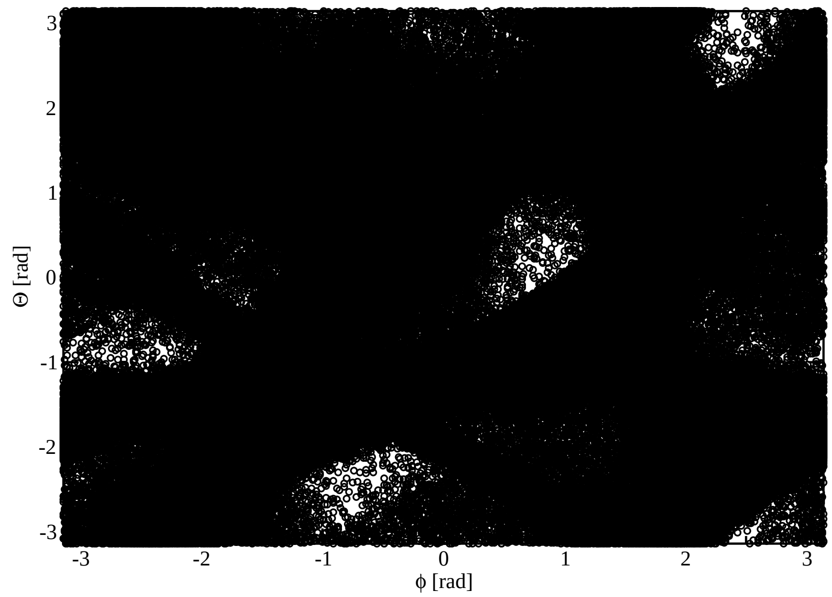


**Fig. 4** Time evolution of the CV during the 975 ns of the WTM simulations.

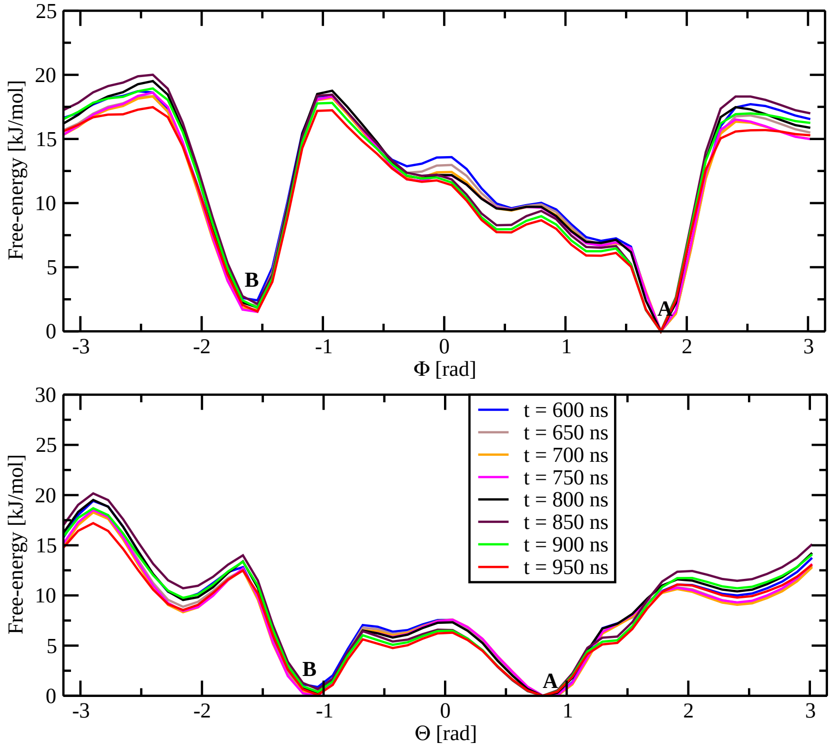


**Fig. 5** Estimates of the integrated free-energy profiles as a function of the WTM CVs deposited along a 700 ns-long WTM simulation time. The selected basins have been shown in Fig. 6 of the main text.

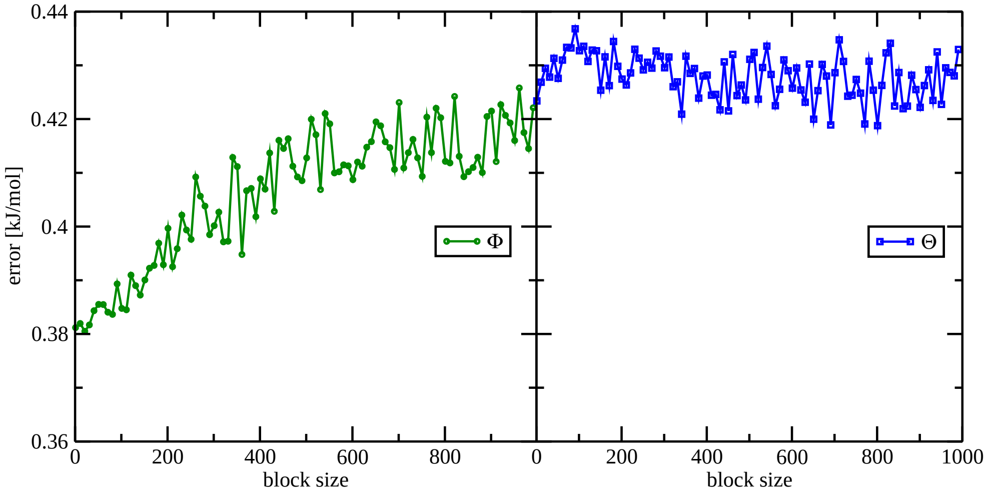


**Fig. 6** Block analysis of the average errors along the free-energy profiles for Θ (left) and Φ (right) as a function of the block length. Taken from the last 400-600 ns of each metadynamics simulation.

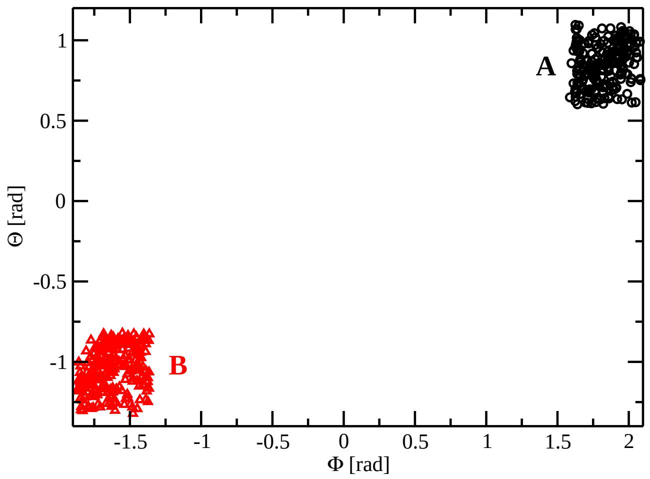


**Fig. 7** Committor analysis of the stable basins (A, B, see Fig.5 of the main text).

### 4 Tables

**Table 1** Aminoacid components of the KRas-4B proteins.

| Full name | Abbreviation |
|---|---|
| Alanine | Ala |
| Arginine | Arg |
| Asparagine | Asn |
| Aspartate | Asp |
| Cysteine | Cys |
| Glutamate | Glu |
| Glutamine | Gln |
| Glycine | Gly |
| Histidine | His |
| Isoleucine | Ile |
| Leucine | Leu |
| Lysine | Lys |
| Methionine | Met |
| Phenylalanine | Phe |
| Proline | Pro |
| Serine | Ser |
| Threonine | Thr |
| Tryptophan | Trp |
| Tyrosine | Tyr |
| Valine | Val |

**Table 2** Parameters employed in metadynamics simulations.

| | |
|---|---|
| Gaussian width of CV1 [rad] | 0.35 |
| Gaussian width of CV2 [rad] | 0.35 |
| Starting (Gaussian) hill [kJ/mol] | 1.0 |
| Deposition stride [ps] | 1 |
| Bias factor | 10 |
| Temperature | 310.0 K |
| Simulation time [ns] | 1000 |

**Table 3** Coordinates of segments forming selected paths for oncogenic KRas-4B-FMe. Along the path, locations of free energy spots (fespot, in kJ/mol) are displayed.

| Minimum free energy path | | | |
|---|---|---|---|
| State | path x | path y | fespot |
| B | -1.61 | -1.07 | 1.00 |
| | -1.70 | -1.26 | 3.51 |
| | -1.63 | -1.50 | 15.80 |
| | -1.51 | -1.71 | 19.21 |
| | -1.30 | -1.85 | 14.09 |
| | -1.07 | -1.77 | 18.61 |
| | -0.84 | -1.65 | 19.39 |
| | -0.61 | -1.56 | 16.68 |
| | -0.36 | -1.48 | 14.33 |
| | -0.13 | -1.35 | 13.84 |
| | 0.09 | -1.21 | 14.13 |
| | 0.31 | -1.07 | 13.32 |
| | 0.54 | -0.93 | 11.96 |
| | 0.77 | -0.78 | 10.55 |
| | 0.98 | -0.60 | 8.33 |
| | 1.18 | -0.40 | 6.61 |
| C | 1.34 | -0.17 | 6.92 |
| | 1.48 | 0.07 | 7.73 |
| | 1.62 | 0.32 | 5.60 |
| | 1.73 | 0.59 | 2.23 |
| A | 1.84 | 0.85 | 0.00 |
| | 1.85 | 0.94 | 0.27 |
| | 1.85 | 1.19 | 6.50 |
| | 1.78 | 1.28 | 14.28 |
| | 1.69 | 1.36 | 21.27 |
| | 1.61 | 1.43 | 27.72 |
| TS | 1.51 | 1.51 | 30.98 |
| | 1.41 | 1.58 | 30.55 |
| | 1.33 | 1.68 | 26.71 |
| | 1.22 | 1.74 | 19.61 |
| | 1.15 | 1.83 | 14.56 |
| | 0.97 | 1.82 | 11.80 |
| | 0.83 | 1.68 | 10.61 |
| | 0.75 | 1.61 | 9.85 |
| | 0.60 | 1.48 | 8.75 |
| D | 0.51 | 1.42 | 8.54 |
| | 0.38 | 1.33 | 9.18 |
| | 0.13 | 1.11 | 12.15 |
| | 0.02 | 0.99 | 12.69 |
| | -0.32 | 0.65 | 12.54 |
| | -0.43 | 0.52 | 13.66 |
| | -0.51 | 0.38 | 14.83 |
| | -0.69 | 0.11 | 16.57 |
| | -0.88 | -0.15 | 18.85 |
| | -0.99 | -0.28 | 19.61 |
| | -1.09 | -0.41 | 18.58 |
| | -1.18 | -0.54 | 14.74 |
| | -1.28 | -0.67 | 9.78 |
| | -1.39 | -0.80 | 4.98 |
| | -1.49 | -0.93 | 1.90 |
| B | -1.59 | -1.05 | 1.00 |

#### Notes and references


1 H. J. Berendsen, D. van der Spoel and R. van Drunen, *Computer Physics Communications*, 1995, **91**, 43–56.

2 H. Lu and J. Martí, *The Journal of Physical Chemistry Letters*, 2020, **11**, 9938–9945.

3 A. Barducci, G. Bussi and M. Parrinello, *Physical review letters*, 2008, **100**, 020603.

4 Y. Zhou, P. Prakash, H. Liang, K.-J. Cho, A. A. Gorfe and J. F. Hancock, *Cell*, 2017, **168**, 239–251.

5 M. Bonomi, D. Branduardi, G. Bussi, C. Camilloni, D. Provasi, P. Raiteri, D. Donadio, F. Marinelli, F. Pietrucci and R. A. e. a. Broglia, *Computer Physics Communications*, 2009, **180**, 1961–1972.

6 G. A. Tribello, M. Bonomi, D. Branduardi, C. Camilloni and G. Bussi, *Computer Physics Communications*, 2014, **185**, 604–613.

7 P. Hošek and V. Spiwok, *Computer Physics Communications*, 2016, **198**, 222–229.
